# Prefrontal brain-to-brain synchrony during human group hunting: Evidence from fNIRS hyperscanning

**DOI:** 10.64898/2026.04.05.716331

**Authors:** E. Yavuz, C. Xu, W. Liu, C. Slinn, A. Mitchell, J. Ali, N. Bloom, N. Khatun, P. A. Kirk, F. E. Zisch, I. Tachtsidis, P. Pinti, F. Ronca, Z. Patai, P. Burgess, A. Hamilton, H. J. Spiers

**Affiliations:** Institute of Behavioural Neuroscience, Department of Experimental Psychology, Division of Psychology and Language Sciences, University College London, London, UK; Emotion and Development Branch, National Institute of Mental Health, Bethesda, Maryland, USA; Bartlett School of Architecture, University College London, UK; Department of Medical Physics and Biomedical Engineering, University College London, UK; Institute of Sport Exercise and Health, University College London, London, UK; Department of Psychology, Ruhr University Bochum, Bochum, Germany; Institute of Cognitive Neuroscience, University College London, UK

**Author notes:** Equal contribution (A. Hamilton, H. J. Spiers).

## Abstract

Orca, wolves, chimpanzees and humans share a similarly impressive capacity for group hunting, where individuals coordinate behaviour together to capture prey. Studying hunting behaviours has important implications for understanding how behaviour in group contexts may be indicative of cognitive decline. Despite growing interest in brain circuits for prey capture, the brain regions involved in tracking prey during a hunt and the behaviours in group hunt linked to success remain unclear. Here we combined functional near infrared spectroscopy (fNIRS) and a virtual minecraft world to examine behaviour, brain dynamics and brain synchrony involved in group hunting behaviour. We focused on the prefrontal cortex (PFC) due to its known role in planning and social coordination and recorded from pairs of individuals as they either cooperated to hunt another person (prey) or simply followed another person. Hunters were more successful if they managed to keep a smaller distance to the prey and moved at speeds that were more synchronised with their co-predator. At high-range frequencies for fNIRS (0.1-0.2Hz), we found greater brain-to-brain synchrony in lateral and medial (frontopolar) PFC regions during hunting compared with chance levels. Together, these findings provide insights into what behaviours and brain dynamics associated with successful group hunting.

## Introduction

Wolves, lions, orca, chimpanzees and humans are among the notable species that show impressive group hunting behaviour (Bailey et al., 2012; Boesch & Boesch, 1989; Pitman & Durban, 2011; Sampaio et al., 2024; Samuni et al., 2018). By working as a coordinated group they significantly increase the efficiency of tracking down and capturing prey. Such collective behaviour is a prime example of a higher order behaviour that requires complex coordination across individuals and planning by each individual to navigate the environment to obtain reward. Size of the group and level of experience of individuals have been found to affect the group performance (Bak-Coleman et al., 2022; Dyer et al., 2008). Despite increased interest in the neural circuits supporting prey capture in solo hunting (Li et al., 2018; Mobbs et al., 2009; Mota-Ortiz et al., 2012; Park et al., 2018), little is known about how brain regions support group hunting, or the process of detecting, tracking, cornering and capturing the prey (Yoo et al., 2021; Goodroe and Spiers, 2022; Chericoni et al., 2026). Studying hunting behaviours has important implications for understanding how behaviour in group contexts may be indicative of cognitive decline (Hamilton et al., 2026).

Key to successful hunting is moving close to the prey to capture it. As prey move to escape and terrain can obstruct direct paths to prey, hunting often involves keeping track of the distance and direction to the prey in order to minimize both for capture. While it remains unclear how brain regions support group hunting, parallels have been noted to navigation, where the distance and direction to a goal must also be minimized in order to arrive at the goal and obtain reward (Goodroe and Spiers, 2022; Spiers, 2022). Mounting evidence indicates a network of brain regions are involved in tracking the distance and direction to goals to support navigation (Spiers and Maguire, 2007a; Viard et al., 2011; Sherill et al. 2013; Howard et al., 2014; Chadwick et al., 2015; Spiers, Olafsdottir and Lever, 2018; Balaguer et al., 2016; Shine et al., 2019; Javadi et al., 2019a; Sarel et al., 2017, 2022; Patai et al., 2019; Bierbrauer et al., 2020; Kunz et al., 2021; Ormond & O’Keefe, 2022; Basu et al., 2021; Liu et al., 2023; Huang et al., 2024). These brain regions include the hippocampus and medial temporal lobe regions, dorsal striatum, retrosplenial cortex, posterior parietal cortex and prefrontal cortex (PFC) (for reviews see: Spiers and Barry, 2015; Epstein et al., 2017; Nyberg et al., 2022). Recent research from human, rodents and bats has revealed that brain regions that code spatial information during navigation can also code spatial information about other agents or moving objects (Danjo et al., 2018; Omer et al., 2018; Polti et al., 2021; Stangl et al., 2020; Wagner et al, 2023) and also for prey capture on a 2-D screen with icons (Yoo et al., 2020; 2021; Chericoni et al 2026).

In humans specifically, a wide range of brain regions (medial temporal lobe, frontal lobe, striatum, retrosplenial cortex and posterior parietal cortex) have been shown to be involved in either retracing the path of a virtual character in a virtual 3D environment (Wagner et al., 2023), predicting the future location of a moving dot on a 2D screen (Politi et al., 2021), or tracking the location of another moving human participant when seated in a real environment (Stangl et al., 2020). However, it remains unclear as to how the human brain keeps track of a moving goal when simultaneously tracking the location of other agents in the environment aside the goal itself, given that these studies did not investigate behaviour or neural activity in a multiplayer context where one is required to coordinate the tracking of multiple moving goals simultaneously in space (Politi et al., 2021; Stangl et al., 2020; Wagner et al., 2023). Studies have also demonstrated that participants gain more reward when foraging with a partner (i.e., in a dyad) than alone in virtual 1D environments, especially when they adapt the frequency of their risk-taking behaviour to match the level of their partners, even in absence of verbal communication (Deng et al., 2025). In real-world environments, humans navigate more successfully in the presence of a stranger than with a friend or alone (Bae et al., 2024). These findings suggest that there may be brain regions which track the location of more than one goal in space simultaneously. However, it remains untested, whether brain regions do track moving goals when navigating with others, i.e., tracking the prey during cooperative hunting.

Similar to hunting, navigation can require taking detours, exploiting opportunistic shortcuts and back-tracking when a well made plan goes awry. Such flexible behaviour during navigation has been associated with the PFC, hippocampus and striatum (Spiers and Maguire 2006a; Spiers and Gilbert, 2015; Javadi et al., 2019b; Gahnstrom and Spiers, 2020; Patai and Spiers, 2021). It seems likely that the PFC would be engaged during group hunting when exploiting the environment to capture the prey.

In addition to evidence of PFC activity tracking the distance and heading direction to a goal in simulated real-world environments (Howard et al., 2014), regions within the PFC play a key role in group coordination and group decision-making (Klein-Fugge et al., 2022; Zoh et al., 2022). The medial PFC is engaged when differentiating social roles, encoding emotional closeness and group belonging (Kieckhaefer et al., 2022; Ron et al., 2022; Roseman et al., 2022), as well as mentalizing (Hill et al., 2017; Jamali et al., 2021; Katsyri et al., 2013; Spiers & Maguire, 2006b), and prospective memory (Burgess et al., 2022). In contrast, the lateral PFC appears important for strategic reasoning and decision-making (Javadi et al., 2017; Nagel et al., 2017; Panidi et al., 2022). Together, lateral and medial PFC appear to support the cognitive and social processes necessary for effective coordination and decision-making in complex tasks, that share similarities with group hunting.

One recent and rapidly developing neuroimaging technique that offers the key advantage for measuring brain activity in two or more people during social interactions or shared tasks - hyperscanning - is functional near-infrared spectroscopy (fNIRS) (Ferrari & Quaresima, 2012). fNIRS uses infrared light (650-950 nm) to monitor changes in the concentration of oxyhemoglobin (HbO2) and deoxyhemoglobin (HbR), providing an indirect measure of neocortical brain activity (Ferrari & Quaresima, 2012). In addition to enabling the simultaneous scanning of multiple brains, fNIRS eliminates the need for participants to remain motionless or restrained, unlike other imaging modalities such as fMRI (Crum et al., 2022; Wilcox & Biondi, 2015). Its portability, improved spatial resolution, and robustness to facial and eye movements, compared to other mobile techniques like EEG, make fNIRS particularly well-suited for studying dynamic, real-world social interactions (Cui et al., 2012; Hakim, 2024; von Luhmann et al., 2021). As such, fNIRS is a well-suited tool for exploring frontal brain activity during group coordination and decision-making in complex tasks like hunting (Crum et al., 2022; Wilcox & Biondi, 2015).

Given the complexity of coordinating as a team in hunting, it seems likely that the hunters’ brains show some degree of real-time neural alignment. Interpersonal neural synchrony (INS) refers to the temporal coordination of concurrent rhythmic brain activities between two or more individuals (Cui 2012; Hakim 2024; Hamilton et al., 2021). In real-world interactions, INS has been associated with behaviour, with INS in the lateral PFC being associated with how competitive players are and their perceived likeability during two-person games (Wen et al., 2025), as well as during group tasks involving more than two players (Yang et al., 2020), whilst INS in the medial PFC has been associated with verbal communication (Nozawa et al., 2016), and with face-to-face deception between two participants (Pinti et al., 2021).

While INS has been linked to various forms of social interaction, it remains unclear how shared neural dynamics relate to successful collective behaviour in tasks that require tracking a moving goal (Nozawa et al., 2016; Pinti et al., 2021). One theory which has been proposed to explain why similar patterns of brain activity are seen between individuals during social interaction and joint tasks is the mutual prediction theory (Hamilton et al., 2021). According to this theory, each person’s brain encodes not only their own behaviour but also predicts their partner’s behaviour. This mutual prediction results in synchrony in brain activity, as the sum of activity in one participant’s brain closely matches the sum of activity in their partner’s brain, reflecting efficient coordination. Other explanations for why similar patterns of brain activity are seen between individuals during social interaction and joint tasks might be due to similarities in task representation, where two or more participants may share similar representations of a given task (Dai et al., 2022). Shared attention (de Felice et al., 2025), and the exchange of information (Nozawa et al., 2016) might also explain why INS is seen during social interaction and joint tasks. Given that hunting requires coordination and decision-making to track group goals, it is plausible that INS in the lateral and medial PFC are linked to group hunting behaviour, reflecting how teams efficiently process shared goals and coordinate their actions in real time.

In this study, we designed a group hunting task in a virtual world created in Minecraft, a sandbox 3D game that supports real-time first-person navigation and interaction between players. We create a world where two human participants hunted another person (the prey) and attempted to catch them in a set time (Hunt trials). We compared brain activity during these Hunt trials with trials in which participants were instructed to follow the prey’s path without the need to capture them (Follow trials). We used fNIRS to investigate brain activity in the PFC during both Hunt and Follow trials. As part of our fNIRS set up we also recorded heart rate, breathing rate and skin conductance, and examined these in relation to trial to trial hunting success (successful prey capture).

We hypothesized several key outcomes. (1) Similar to navigation (e.g. Howard et al., 2014) the distance and direction to the prey would positively correlate with activity in PFC regions during the Hunt trials (increasing the greater the distance and greater the angle needed to turn towards the prey). (2) This correlation would be greater during Hunt trials compared with Follow trials. This was predicted on the basis that, while the Follow trials were visually and motorically similar to Hunt trials, in Hunt trials there should be increased demand to monitor the distance and direction to the prey for capture. This prediction is based on evidence of increased PFC tracking of distance and direction to the goal previously reported in active goal tracking (navigation) compared to following guidance instruction on which directions to travel (Howard et al., 2014; Patai et al., 2019). (3) Hunt trials, where two predators actively coordinate to hunt prey together, would show greater PFC synchrony than Follow trials, due to the greater demand to coordinate behaviour, parallelling prior fNIRS studies of cooperative behaviour (Dai et al., 2022; Pinti et al., 2021; Nozawa et al., 2016; Yang et al., 2020).

In addition to the hypothesis-led fNIRS predictions we also explored how measures of behaviour, such as the distance between the two predators or variability in orientation towards the prey, and physiological measures (heart rate, breathing rate and skin conductance) might be related to hunting success. Such behavioural measures were also entered into exploratory analysis of fNIRS data.

## Methods

### Participants

Participants aged 18-45 years old were recruited through social media platforms including LinkedIn and Instagram, as well as through the recruitment platform SONA. Inclusion criteria required participants to be physically healthy, with normal or corrected-to-normal vision, and without any neurological or psychiatric disorders. Participants were also required to have at least some experience playing video games using keyboard controls. The experimental procedure conformed to relevant guidelines and regulations, and received ethical approval from the research ethics board at UCL conforming to Helsinki’s Declaration (EP 5975/003). Participants were reimbursed £10 per hour for their participation. In total, 32 participants were recruited (14 men, 18 women; mean age = 21.5 years, SD = 3.57 years, range = 18–33 years). Eight participants were excluded due to poor fNIRS data quality, primarily caused by motion artifacts, as well as either insufficient signal intensity (light penetration) or excessive light penetration. This resulted in a final sample of 24 participants (11 men, 13 women (3 male-male pairs, 5 female-female pairs, 4 female-male pairs), mean age = 21.1 years, SD = 3.35 years, range = 18–33 years).

### Sample size calculation

As this study was exploratory, we did not calculate a desired sample size a priori. However, other studies looking at the neural basis of navigation in virtual environments have used similar sample sizes (Howard et al., 2014; Patai et al., 2019). As mentioned above, the final sample size used for neural analysis is lower than that used for behaviour, given lost data.

### Experimental Task

#### Environment

A virtual environment was constructed in Minecraft, a sandbox video game created by Mojang Studios and owned by Microsoft (Figure 1). It allows first-person-view navigation of virtual worlds, easy creation of custom environments and the extraction of positional data from the game (Stark et al., 2021). The Minecraft world featured boulders and rivers dividing the world into multiple islands connected by bridges (Figure 1). Boulders intermittently obscured the prey, while rivers and bridges increased the need for spatial planning in order to capture the prey. The large environment was intended to encourage spatial planning and simulation of the prey’s future trajectories. Soul sand, a Minecraft block which slows players down when walked on, was placed on the bridges to slow players down. Prey entering a bridge would slow down allowing hunters to catch up, and then when off the bridge allow them to escape faster. This helped increase the variation in distance to the prey over the course of trials, which otherwise would tend to vary little once the predators were close to the prey. Landmarks, including a central orange tower and a repeating pattern of orange and blue blocks at opposite sides of the world, were added to enhance spatial orientation.

**Figure 1.**
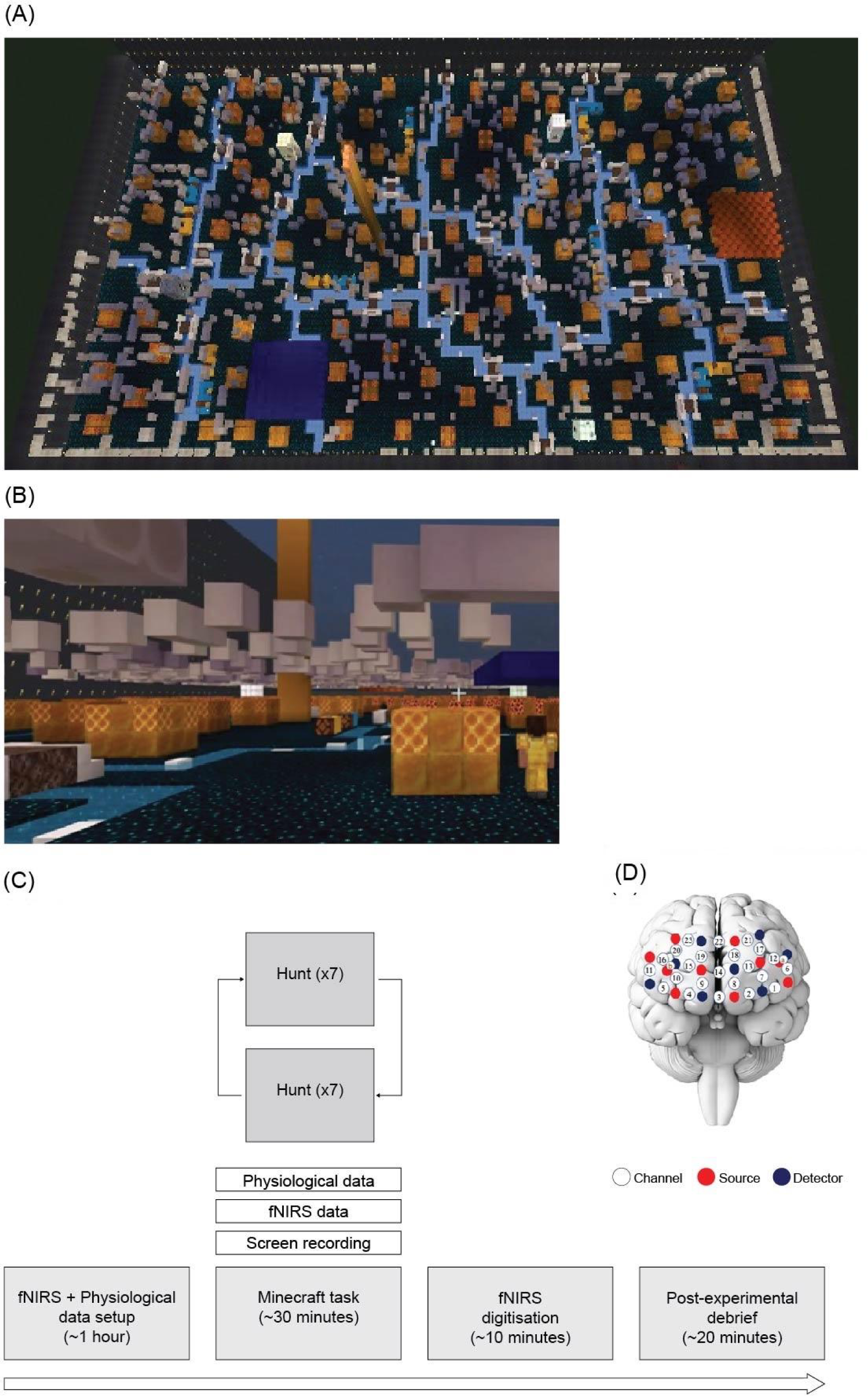
Minecraft world and experimental design. **(A)** Top-down view of the Minecraft world, showing rivers that separated the world into multiple islands connected by bridges. Soul sand on the bridges slowed the players down, creating variation in Euclidean distance to prey and increasing the hunting challenge. The central orange tower served as a primary landmark, a repeating series of orange blocks near the right (east) wall and blue blocks near the bottom-left (south-west) wall also functioned as landmarks to aid orientation. Participants never saw this view, and experienced the environment in first-person at ground level. **(B)** First-person view from a predator’s perspective during a Hunt trial. Prey is visible on the right as the character in yellow armour. **(C)** Trials pseudorandomly alternated between Hunt and Follow trial types with no more than two in a row of the same type. The length of the Hunt trials varied based on the predators’ success in hunting the prey. Follow trials were pseudo-randomly selected to last between 30 and 60 seconds to approximate the average length of the Follow trials. Functional Near-Infrared Spectroscopy (fNIRS) and physiological data (heat rate, breathing rate, GSR) were recorded during the task but not during the post-experimental debrief. **(D)** A wireless fNIRS system from Artinis was used to collect neural data from the two predators. Each headset consists of 10 light sources (red) and 8 detectors (blue), creating 25 measurement channels, which were placed on the scalp region covering PFC.

### Task

Before starting the main task, participants had 5 minutes to explore the Minecraft world to ensure they were comfortable with navigating the world and were familiar with its layout. After completing this exploration phase, participants started the main task. Participants did not see a map of the environment before starting the task, and thus had to learn the structure of the environment through exploration.

Due to the limited period in which fNIRS can be worn with comfort, 7 Hunt and 7 Follow trials were presented to participants in the main task, taking approximately 40 minutes in total. In each trial two recruited participants took on the roles of predators (working cooperatively), while the prey was a confederate (a member of the research team), all logged into the same Minecraft server. Different members of the research team acted as the prey in different sessions. All were experienced over many sessions with acting as the prey.

In Hunt trials, the two predators had to chase after and slay the prey with a virtual diamond sword before 60 sec had elapsed. If either of the predators managed to kill the prey before the end of the trial, the predators would score a point for the team. Failure meant no point was scored and the prey would score a point. Because the prey could outrun the predators, success in this task required predators choosing future paths that would intersect where their prey might be in the future. By contrast in Follow trials, the predators simply had to follow in the path of the prey as closely as possible, e.g. if the prey proceeded forward then turned left, they proceeded to the point it turned left and then followed its path after that point. Thus, for Follow trials there was no need to outmanoeuvre the prey, simply to follow a path. There were no points awarded in Follow trials and no requirement to slay the prey. The order of Hunt and Follow trials was pseudorandom with no more than 2 trials of one type in a row.

At the start of each trial, the predators were teleported to one of seven pseudo-randomly chosen locations, with the two predators always starting each trial on the same side of the river, but with the prey on the opposite side of the river. This initial starting position was to prevent the predators from catching the prey immediately, encouraging them to engage in spatial planning to effectively locate and pursue the prey. No start location would appear more than once for each trial type, preventing the use of habitual or learned strategies when hunting the prey. At the start of each trial, a message would appear on screen telling the predators which trial they were about to complete (Hunt or Follow), followed by a message to tell the predators to begin the trial. This latter message was accompanied by a light turning on to indicate the start of the trial. The prey was dressed in blue in Hunt trials and gold in Follow trials in order to help the predators distinguish between trial types whilst in the tasks. Based on pilot data collection the length of Follow trials was selected to range between 30 and 60 seconds to match the anticipated length of the Hunt trials. Hunt and Follow trials lasted on average 42.8 and 42.6 seconds, respectively. During each trial, participants were instructed not to sprint or jump so that speed was controlled by other features of the world. Their speed was set to a minimum of 4.317 m/s, which is the default walking speed in Minecraft (Minecraft Wiki, 2023a).

### Experimental procedure

The participants sat in their own booth in the experimental room, each containing a laptop on which they would complete the task (Figure 1C). Each booth was separated by a divider, meaning the two participants could not see each other during the task. Participants were instructed not to speak to each other during the task for several reasons. Firstly, it enables this study to be replicated using other imaging modalities such as fMRI, where participants are unable to speak to one another. Secondly, in many commercial multiplayer video games, participants do not communicate verbally (McLaren-Gradinaru et al., 2023). Thirdly, it reduces breathing-related artefacts in the fNIRS signal. The prey sat in their own booth with a provided laptop, which was separated from the predators’ booths, ensuring that neither of the predators could see the prey and vice versa. After consent to the study participants received instructions on how to wear the equipment for measuring physiological signals (heart rate, breathing, and skin conductance).

After putting on the equipment, participants reviewed the task instructions and were informed that they would not be allowed to communicate during the task, and that their laptop screens displaying minecraft would be recorded for later analysis. After setting up the fNIRS system participants completed two practice trials (e.g. 2 hunt, 2 follow) as needed before beginning the main task.

After completing the task, participants underwent a digitisation session using the Polhemus Patriot System, PiMgr software (version 2.7.3, Polhemus), where the location of fNIRS channels on their headset was recorded to ensure that the anatomical locations of channels were consistent across participants. Finally, participants underwent an experimental debrief with a retrospective verbal report collected (see Spiers and Maguire, 2006b; 2007b; 2008; Gregorians et al., 2025 for similar protocols). During this process participants individually watched a video replay of their performance in hunting task trials. While watching they were asked to verbally report any strategies they remembered using (e.g. “I thought I would head to the left as that is the shortest path a bridge to get to the prey”) as well any recollections of their thoughts and feelings (e.g. “When the prey suddenly came into view I was surprised and thought I realised I need to outwit them by taking a right round this set of blocks”). Quotes here are illustrative. A member of the research team coded their responses numerically on a second-by-second basis, each second was assigned 0 for each second when the participant provided no report on a particular thought or feeling, and 1 to indicate when they did, and which predetermined category this came from. Categories were determined from pilot data and included the following: surprise, thinking about the other predator or the prey, spatial planning, expectation confirmation, happiness, and frustration. The entire experimental procedure took approximately 2 hours to complete, including setting up the fNIRS (Figure 1).

### Behavioural, neural and physiological data acquisition

#### Behavioural data

The Minecraft game was hosted on an external server provided by Shockbyte (Multicraft - Home, 2023), using the Paper software (version 1.20.2) which allowed for improved server performance and enhanced gameplay. Custom scripts to extract various behavioural variables from Minecraft including positional information and timestamps were created using the Java programming language (version 17) within the Eclipse Integrated Development Environment (version 4.25.0) using Apache Maven (version 3.10.1), a widely used tool for project management in Java development. Using these custom scripts, behavioural data was collected whilst participants completed the experiment, recorded at a 1 Hz frequency.

### Physiological data

Skin conductance (EDA), heart rate (ECG) and breathing rate data was collected using the biopac system, at a sampling frequency of 2000Hz.

### fNIRS data

A wireless fNIRS system from Artinis was used to collect neural data, consisting of 2 headsets. Each headset consists of 10 light sources and 8 detectors, creating 25 measurement channels, which were placed over the scalp to record PFC regions (see Figure 1). The light sources were emitting near-infrared light at 753 nm and 844 nm. The sampling frequency of the system is 25 Hz. To place the headset in a reliable manner across both participants, we used the 10-20 electrode placement system and aligned channel 3 to the Fpz-Fz line. A 3D magnetic digitizer (Fastrak, Polhemus) was then used to digitise the locations of the optodes (i.e., sources and detectors) and the 25 channels and 5 landmarks (Nasion, Inion, right and left pre-auricular points and Cz). The digitized coordinates were converted into the MNI space and co-registered onto a standard brain template using the NIRS SPM software package (Ye et al., 2009). The median MNI coordinates of the 25 channels across all participants and the corresponding anatomical location was then determined according to the Broadmann Areas atlas (Zhang et al., 2017) (Table 1).

**Table 1.**
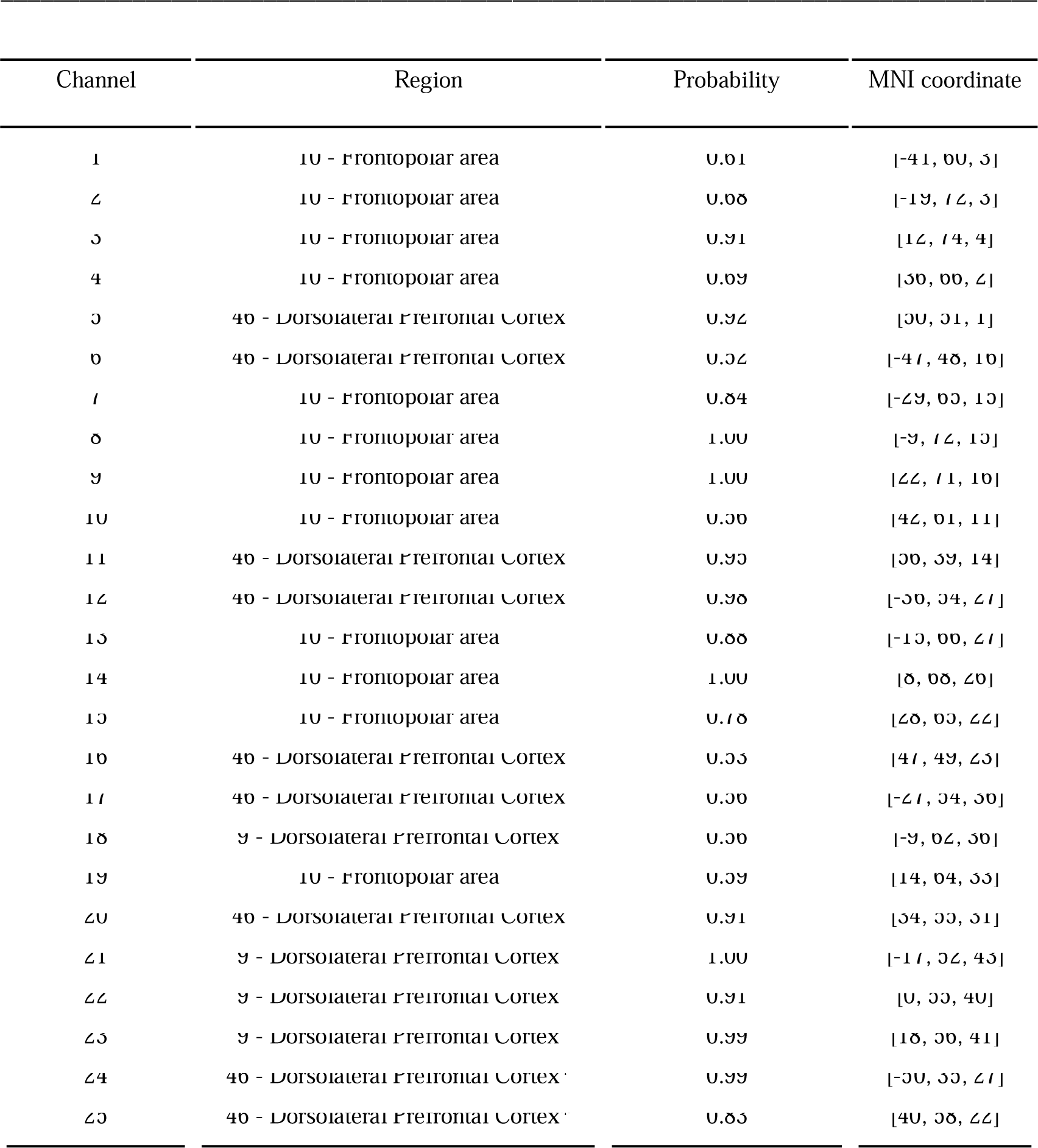
fNIRS channel locations. Regions of interest corresponding to each fNIRS channel were approximated using the Brodmann Atlas. * = Channels 24 and 25 are classified as short channels, designed to exclusively capture extracerebral signals, such as blood pressure waves, Mayer waves, respiration, and cardiac cycles. The signal components obtained from these short separation channels can be considered as “noise” in the signals recorded from long channels (Brigadoi & Cooper, 2015).

### Data preprocessing

#### Behavioural data

Behavioural data was extracted using scripts written in Java and then pre-processed to ensure reliability. This process involved removing duplicate timepoints and linearly interpolating missing ones due to missing time points in the Minecraft server. We also removed data at points where participants accidently ran or jumped during the task, linearly interpolating over the missing data points. Such missing timepoints were infrequent, occurring approximately 5 times across sessions. We calculated several behavioural metrics derived from the x, y, and z coordinates of each predator (designated as ‘self’ and ‘other’) to analyse interactions between the self, the other predator and the prey. These metrics were divided into second-by-second and trial-based measures. For the second-by-second metrics, we first computed the Euclidean distance between the self and the prey, as well as between the self and the other predator. Additionally, we calculated the heading direction from the self to the prey. This was defined as the angular difference between the self’s facing direction and the direction from the self to the prey. To ensure that left and right deviations contributed equally, angular differences were transformed from the 0–360° range to 0–180° (i.e., 360°–θ for angles >180°), following the approach described in Howard et al. (2014). This provided a final value between 0 and 1, where a value of 0 indicated that the self’s facing direction was perfectly aligned with the direction from the self to the prey, whilst a value of 1 indicated the opposite (i.e. misalignment). We also calculated the heading from the self to the other predator, which was calculated in the same way as described for the heading direction from the self to the prey (See Figure 2).

**Figure 2.**
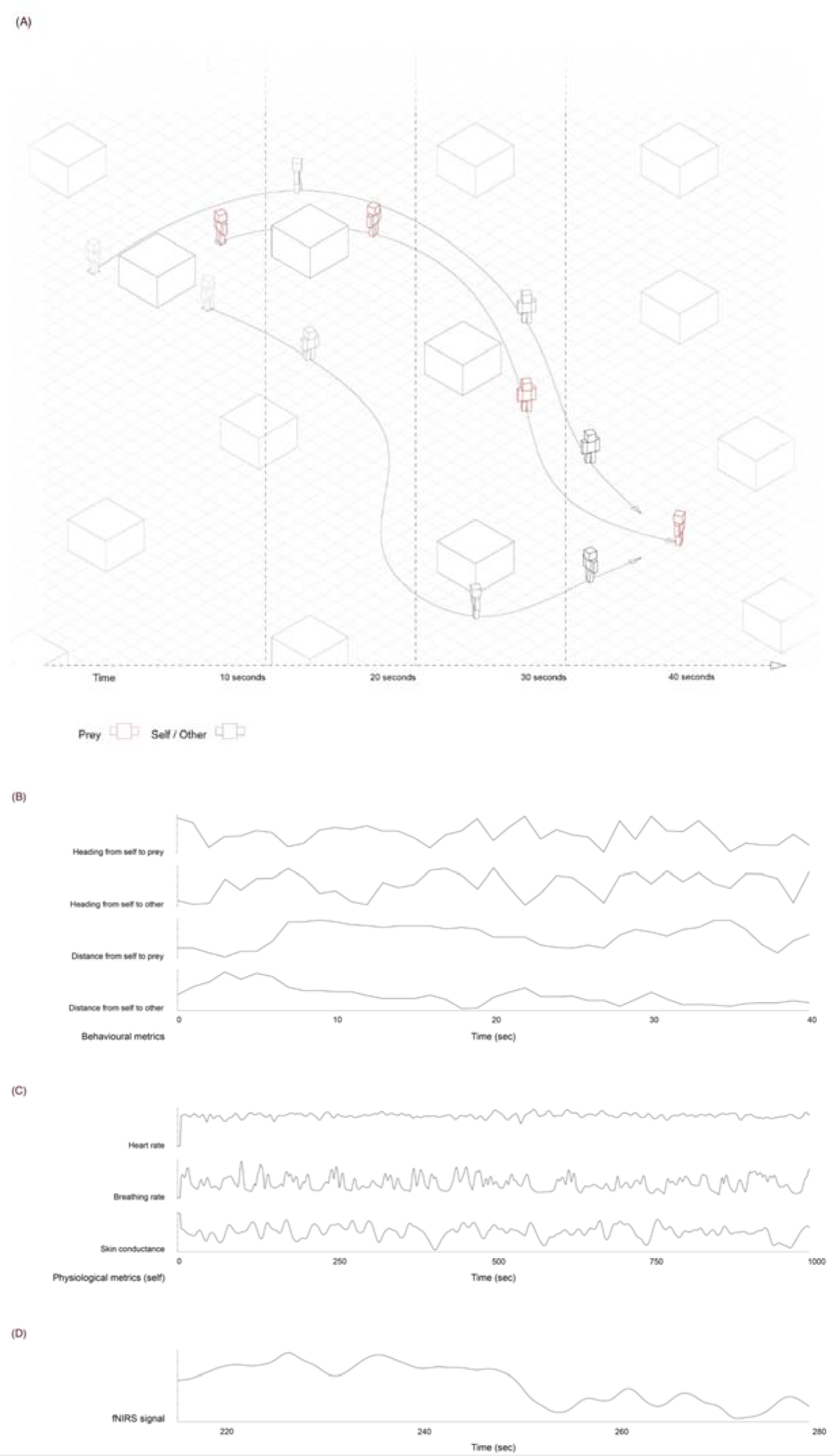
Behavioural, physiological, and fNIRS data collected during each trial. **(A)** Trajectories of the self, the other predator and the prey during an example Hunt trial at each point in time, overlaid onto a top-down schematic of the Minecraft world. Behavioural metrics include the Euclidean distance between self and the prey, the Euclidean distance between the self and the other predator, the heading direction from self to the prey, and the heading direction from the self to the other predator. See Movie1 and Movie2 for plots of the trajectory of the participants over time in hunt and a follow trial. **(B-D)** Behavioural **(B),** physiological **(C)**, and neural data **(D)** collected during an example Hunt trial. Physiological metrics include the heart rate, breathing rate, and tonic skin conductance of the self, shown here as an example. The data were collected and analysed for both the self and the other predator. **(D)** An example from one of the 25 fNIRS channels is shown for illustrative purposes, though the data was collected and analysed for all channels.

For trial-based metrics, we calculated the synchrony of the self and the other predators speeds by computing the correlation between their second-by-second changes in path distance travelled. If both predators were moving toward the prey at similar rates, their movements would be positively correlated. In contrast, if one predator was pursuing the prey while the other took a divergent route, their movements would be less correlated or even decorrelated. To capture how this synchrony changed over time, we used a rolling 5-second window to compute correlation values, and then averaged these values across the trial, yielding a measure of the synchrony of the predators speeds across the trial. We refer to this resulting measure as the speed synchrony between predators.

To ensure the reliability of our results, we checked for model assumptions and identified multivariate outliers using Mahalanobis’ Distance. This method has been shown to have high sensitivity and specificity, with minimal bias compared to other techniques when detecting multivariate outliers in simulated and real datasets (Curran, 2016; Ward & Meade, 2022; Zijlstra et al., 2011). In total, 23 data points (hunt trials that lasted for very short durations) were identified as outliers and removed from the analysis, while no participants were excluded. The results presented here did not differ when we included the excluded data points.

### Physiological data

Heart rate, breathing rate, and skin conductance data were preprocessed using the Python packages neurokit2 (Makowski et al., 2021) and RapidHRV (Kirk et al., 2022). After preprocessing, data was downsampled to 1 Hz to align with the sampling frequency of the behavioural and fNIRS data for analysis. Thus, the final set of physiological variables included heart rate (BPM), breathing rate, and tonic skin conductance.

### fNIRS data

Concentration changes in oxy- and deoxy-Hb were calculated by the Artinis system using the modified Beer-Lambert law. Raw HbO2 and HHb data were down-sampled to 1 Hz and motion artefacts were identified and corrected using the wavelet-based approach (Molavi & Dumont, 2012) implemented by the Homer2 software package (Huppert et al., 2009). Physiological noise (e.g., heart rate and breathing rate) and slow trends in the signal were removed using a Butterworth band-pass filter between 0.01 and 0.4 Hz. The correlation-based signal improvement (CBSI) preprocessing step (Cui et al., 2010) was then used to combine preprocessed HbO2 and HHb signals into an ‘activation signal’ (Scholkmann et al., 2014). The CBSI method was applied for two reasons: 1. It allows one to infer functional brain activity based on only one signal which includes information on both HbO2 and HHb and 2. It increases the contrast-to-noise ratio of fNIRS signals, reducing the impact of non-neuronal systemic confounding factors, such as extracerebral signals from blood pressure waves, Mayer waves, respiration, and cardiac cycles. The activation signal for each of the 25 channels over the experimental conditions was then fit to an individual design matrix for each participant using a Matlab script calling the estimation functions from NIRS-SPM (Ye et al., 2009).

### Data Analysis

#### Behavioural and physiological data

We first conducted an exploratory analysis to examine the relationships between behavioural and physiological metrics and their associations with hunting success. Based on this, we conducted a linear mixed-effects regression model, using the time taken to catch prey as the primary outcome of hunting success. Although we also calculated the proportion of successful trials, these two variables were highly correlated. We chose the time variable for its continuous nature, compared to the binary success rate. To address multicollinearity, we calculated the variance inflation factor (VIF) for each variable, considering VIF values below 5 acceptable (Fox & Monet, 1992; Kim, 2019).

The model incorporated several metrics (predictors), where for each predictor, an average value was calculated per trial, resulting in a single value representing the overall behaviour for that metric during each trial. All predictor variables were standardized by z-scoring prior to inclusion in the analysis. The predictor variables were the distance between the self and the prey, the distance between predators, the heading direction from the self to the prey, the heading direction from the self to the prey, the heading direction from the self to the other predator, the speed synchrony between predators, the heart rate of the self, the breathing rate of the self, and time. Participants and dyad were treated as random effects to address variations in performance between dyads and among participants within each dyad. We also conducted an additional linear mixed effects model to examine the association between the debrief data and hunting success. The variation in distance and direction was similar across hunt and follow trials (*p* > .05).

As an exploratory analysis, to test for leader–follower dynamics, we applied the pull event detection method recently used in another human foraging task (Wu et al., 2025). Briefly, candidate pull events are defined as *min–max–min* sequences in pairwise distance and are filtered by strength (change in distance relative to absolute distance), disparity (asymmetric movement between participants), leadership (role reversal across segments), and duration (≥3 s). For implementation details, see Wu et al. (2025).

### Neural data

#### Single-Subject Analysis

The input variable for the first-level analysis of the fNIRS data was the Correlation Based Signal Index (CBSI), reflecting brain activation during the task. We used 6 different design matrices to explore different regressors as summarised in Table A1. Design Matrix 1 modelled the Euclidean distance and the heading direction from the self to the prey separately for Hunt and Follow trials, as well as heart rate, breathing rate, activity unrelated to the task, and time across the experiment, to account for potential confounding factors.

To test our hypothesis that activity in the lateral (lPFC) and medial prefrontal cortex (mPFC) would relate to the Euclidean distance and the heading direction from the self to the prey, we conducted one-sample t-tests on the beta values from the Hunt and Follow trials to determine if neural representation was greater than chance in each condition. We then performed a paired t-test to compare the neural representation of the Euclidean distance and the heading direction from the self to the prey between the Hunt and Follow trials. Additionally, we examined the reverse association to assess whether activity was stronger during Follow trials, interpreting the t-statistic direction accordingly (Design Matrix 1). False Discovery Rate (FDR) was used to correct for multiple comparisons across channels. The p-values obtained from the statistical tests for each channel were ranked, and the FDR procedure was applied using an alpha threshold of .05. Specifically, for each p-value, we compared it to the threshold *i*/25 x .05, where *i* is the rank of the p-value and 25 is the total number of comparisons. The largest p-value that satisfied this threshold was considered significant, and all smaller p-values were also considered significant. The same method for calculating FDR was applied consistently across all analyses.

As part of an exploratory analysis, Design Matrix 2 (Table A1) modelled Hunt and Follow trials as task regressors, along with the confounds of heart rate, breathing rate, activity unrelated to the task, and time, to examine whether brain activity differed significantly between Hunt and Follow trials. We conducted linear mixed effects models to assess whether beta values from the Hunt trials and the Hunt > Follow contrast were associated with hunting success. FDR was used to correct for multiple comparisons across channels.

Additionally, Design Matrices 3–6 were included as part of an exploratory analysis and modelled the following: heading direction to the prey and to the other predator independently (Design Matrix 3); distance to the prey and to the other predator independently (Design Matrix 4); the interaction between the heading direction from the self to the prey and the Euclidean distance to the prey (Design Matrix 5); and the speed synchrony between predators (Design Matrix 6).

All matrices also included heart rate, breathing rate, activity unrelated to the task, and time as confounding factors. We conducted one-sample t-tests on the beta values for the Hunt > Follow and Follow > Hunt contrasts across all design matrices to determine if activity was significantly greater than chance in each condition. FDR was used to correct for multiple comparisons across channels.

Finally, we investigated whether participants’ verbal descriptions of their thoughts and feelings during the Hunt trials (from the debrief data) were related to activity in those same trials. We categorised their verbal descriptions into six cognitive processes: surprise, thinking about the other predator or the prey, spatial planning, expectation confirmation, happiness, and frustration. For each participant, we calculated a single score per trial representing the proportion of the trial spent experiencing each cognitive process, and then summed these scores across trials to produce one score per participant. Given that participants’ ratings for individual cognitive processes were highly multicollinear across categories, we derived two summary measures: the average score across all categories and the first principal component from a principal component analysis of these ratings. Linear mixed effects models were then conducted to assess the relationship between these two summary cores and the corresponding beta values from the Hunt trials and the Hunt > Follow contrast, using Design Matrix 3. To ensure that any associations were not an artifact of the fact that some participants reported a greater number of cognitive processes than others, we also conducted a model where we included the total number of reported events for each participant as a predictor variable. FDR was used to correct for multiple comparisons across channels.

### Dyadic Analysis

Although the single-subject analysis had a final sample of 24 participants, the final sample for hyperscanning consisted of only 18 participants (9 dyads) given that when participants were excluded due to poor fNIRS data quality, primarily caused by motion artifacts, as well as either insufficient signal intensity (light penetration) or excessive light penetration, their partners also had to be excluded from the dyad.

Among the majority of fNIRS-based hyperscanning studies, wavelet transform coherence (WTC) has been the most commonly used to assess INS (i.e. measure the strength of brain-to-brain synchronisation) during social interactions (Cui et al., 2012). INS between the self and the other predator was assessed using the wavelet coherence method, which measures the alignment of neural signals between participants over time and across different frequency bands (Cui et al., 2012; Hakim et al., 2023). This approach captures the dynamic nature of inter-brain synchrony during joint hunting tasks. The analysis was performed on a channel-wise basis to assess synchrony across different brain regions.

To test the predictions that actively hunting prey alongside another predator (Hunt trials) would result in greater brain synchronisation than chance levels, and that passively following an experimenter with another predator (Follow trials) would result in greater brain synchrony than chance levels, we employed a time-scrambling method. For each channel and dyad, the raw data - containing data from both Hunt and Follow trials - were time-shifted 1,000 times to create 1000 datasets where the phase of the wavelet coherence data is kept intact but where the neural signal of one participant is shifted relative to their dyad partner. This effectively breaks the temporal alignment between participants, controlling for any synchrony that could arise simply due to the timing of activity rather than true interaction. To assess whether observed coherence values were significantly greater than expected by chance, we compared them to the null distribution of WTC values generated from time-scrambled data. This meant that for each channel in each dyad, we obtained a statistic telling us if the observed WTC falls in the top 5% or 1% of the null distribution of WTC values for that channel. Given that our dataset contained 25 channels in 9 dyads, we wanted to obtain a whole-array p-value which told us whether the activation of a particular channel in all these dyads was likely to arise by chance.

Following the logic of group-level statistics in fMRI analysis in SPM, we considered two thresholds – the *alpha-threshold* that applies to each individual channel, and the *k-threshold* which defines the number of dyads which must show an effect. To choose appropriate values for these thresholds, we used Monte Carlo simulation to define if an effect could arise by chance. For 9 dyads, the k-threshold could range from 1 to 9, and for each k-threshold value, we used the simulations to estimate the corresponding *alpha threshold* which ensures a false discovery rate (FDR) of .05 across the entire set of channels. For the primary analysis, we used a k-threshold of 4, corresponding to an alpha threshold of 0.180. Channels meeting this criterion were deemed significant. We verified that the results were robust by testing k-thresholds of 3 and 5, which yielded consistent findings.

For all analyses, we calculated the average wavelet coherence in the low (0.02-0.03 Hz), medium (0.03-0.1 Hz), and high (0.1-0.2 Hz) frequency bands for each channel and dyad during both Hunt and Follow trials, which represented INS during these conditions. The low frequency band consists cycles lasting ∼33-50 seconds, representing slow, sustained processes that likely span the length of the whole trial. The medium consists of cycles lasting ∼10-33 seconds, representing events that likely unfold over substantial portions of a trial. The high frequency band consists of cycles lasting ∼5-10 seconds, representing faster cognitive processes (De Felice et al., 2024). Using these frequency bands allow INS to be examined across multiple temporal scales, ranging from slow, sustained alignment to faster, interaction-driven coordination, while remaining within the physiological and technical constraints of fNIRS; a time period of less than 5 seconds (frequencies >0.2 Hz) is unmeasurable given that the hemodynamic response function takes 5 seconds to peak, frequencies <0.01 Hz are indicative of baseline drift, and frequencies ∼1Hz are indicative of cardiac signals and motion artifacts (Chen et al., 2025). Given the time-pressured nature of the task, we chose to focus on the medium and higher frequency bands, given that continuous action in response to the prey would be required to successfully catch the prey and be more likely to occur over a shorter timescales, although we also explored neural responses in the low frequency band.

We also conducted two contrasts to directly compare Hunt and Follow trials. First, we performed a Coherence Hunt > Follow contrast by subtracting the average coherence of Follow trials from that of Hunt trials, followed by a one-sample t-test to determine if synchrony was significantly greater during Hunt trials. Second, we conducted a Coherence Follow > Hunt contrast, subtracting the coherence of Hunt trials from Follow trials, and again used a one-sample t-test to assess whether synchrony was greater during Follow trials. Unlike the primary analysis, which used a Monte Carlo–based procedure to account for multiple comparisons across channels and dyads, these contrasts were corrected using a standard FDR procedure across channels.

Additionally, we calculated the proportion of time in which both members of a dyad reported experiencing the same cognitive processes, producing a single score per trial, and then summed these to produce an average score per dyad. We conducted a linear mixed effects model to determine whether these scores were associated with the coherence values from the Hunt > Follow contrasts, as well as from the Hunt and Follow trials separately. To ensure that any associations were not an artifact of the fact that some participants reported a greater number of cognitive processes than others, we also conducted a model where we included the average number of reported events for each dyad as a predictor variable. FDR was used to correct for multiple comparisons across channels.

Both the preprocessing and generation of design matrices were performed in MATLAB (version 22b), while statistical analyses and visualizations were conducted using R (R studio version 2024.02.2+764, R version 4.5.1), and Python (version 3.9.12).

## Results

### Behavioural

We first investigated the behavioural variables associated with successful hunting. For the self-to-prey metrics, the Euclidean distance between the predator and the prey was significantly positively associated with the time taken to catch the prey (β = 3.24, CI = [1.29, 3.24], t = 3.26, *p* = 0.007, f² = 0.92). These results suggest that predators were more successful when they stayed closer to the prey.

Speed synchrony between predators was significantly negatively associated with the time taken to catch the prey (β = -13.13, CI = [-20.88, -13.13], t = -3.32, *p* = 0.007, f² = 0.97; Figure 3), suggesting that the more synchronised the speeds of the predators were, the faster they were able to catch the prey.

**Figure 3.**
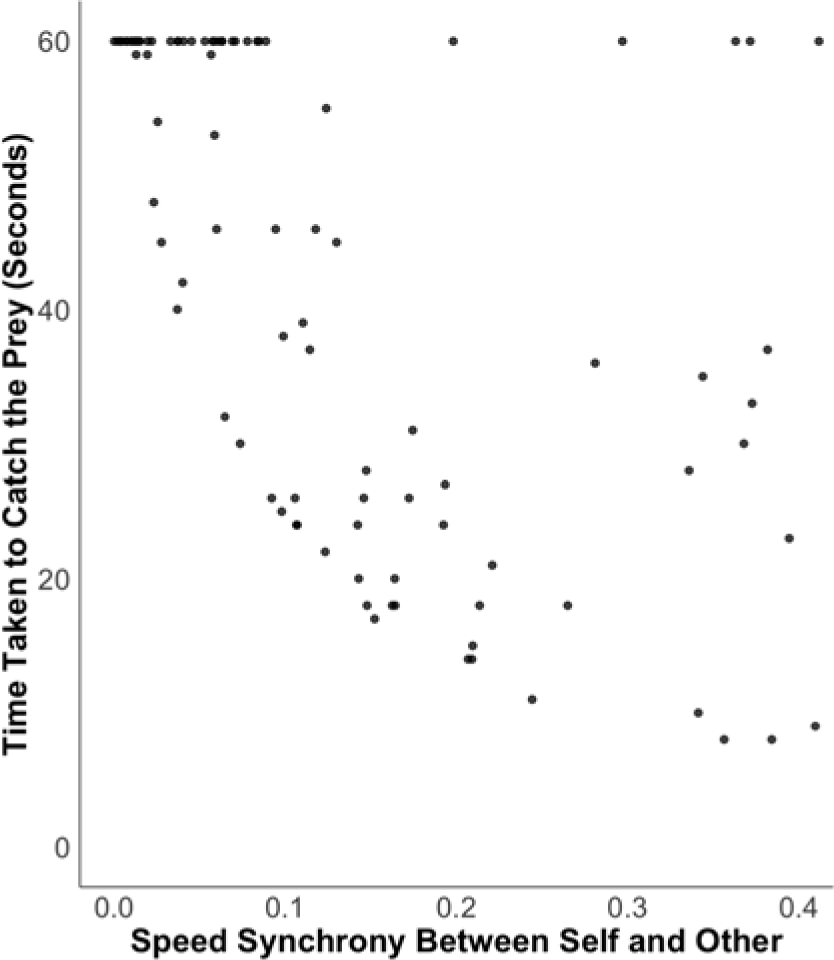
Hunting success (quick capture) is associated with greater synchronisation of the speed of the predators. Speed synchrony between self and other is plotted against the time taken to catch the prey, with each dot indicating a dyad, for each trial in the experiment. The shorter the time, the more successful the hunt. When synchrony in speed between predators is higher, hunting success is higher. Although other variables are also significant in this model, only this variable showed an effect size above 0.1. Each data point represents the score for a single dyad on a given hunt trial. 60 seconds was the maximum limit on the trial.

Other behavioural variables, distance to the other predator, heading direction to the prey, heading direction to the other predator, heart rate, breathing rate, and time across the experiment were not significantly related to success (all *p* > .05, f2 < 0.15). Please see Appendix Figure S1 for videos of the trajectories of the predators and the prey during Hunt and Follow trials.

In sum, successful hunting (rapidly catching the prey) was associated with maintaining closer proximity to the prey and greater speed synchrony between the predators. No significant effects were found for physiological measures or elapsed experimental time. An association between the distance to the prey and the time taken to catch the prey is unsurprising given the successful trials end with close proximity. Other variables are less easily explained and will require replication.

**Table 2.** Model predicting the time taken to catch the prey during Hunt trials based on behavioural and physiological metrics. P-values for the significant associations are highlighted in bold.

| Variable (average across trial) | $\beta$ | 95% CI | $t$ | $p$ | sig | $f^2$ |
| --- | --- | --- | --- | --- | --- | --- |
| Distance between self and prey | 3.24 | [1.29, 3.24] | 3.26 | **0.007** | * | 0.92 |
| Distance between predators | -0.12 | [-5.46, -0.12] | -0.04 | 0.965 |  | <0.01 |
| Heading direction from the self to the prey | 1.32 | [-0.58, 1.32] | 1.36 | 0.182 |  | 0.06 |
| Heading direction from the self to the other predator | 0.18 | [-1.78, 0.18] | 0.18 | 0.861 |  | <0.01 |
| Speed synchrony between predators | -13.13 | [-20.88, -13.13] | -3.32 | **0.007** | * | 0.97 |
| Heart rate self | 0.19 | [-1.71, 0.19] | 0.20 | 0.846 |  | <0.01 |
| Breathing rate self | 1.18 | [-1.02, 1.18] | 1.05 | 0.314 |  | 0.10 |
| Time across experiment | -1.12 | [-2.73, -1.12] | -1.36 | 0.177 |  | 0.01 |

### Single-subject Brain Activity Associations with Distance and Direction

When correcting for multiple comparisons across channels, we found no significant association between brain activity and the distance or direction to the prey or other predator in either the hunt or follow trials, or when Hunt and Follow trials were combined to a single regressor (*p* > .05). At a more liberal uncorrected threshold, we did find a positive association between activity in channel 18 (over left dlPFC) and the Euclidean distance to the prey (*p* = 0.035) when Hunt and Follow trials were combined to a single trial regressor (Table 3). No other associations were evident.

**Table 3.** Channels with uncorrected significant correlations with spatial metrics. Only significant results (*p* < .05) are shown. Numbers indicate channel numbers (15 = channel 15). Direction is the egocentric direction, collapsed over left and right where 0 = ahead and 1= behind. Distance is the Euclidean distance where 0 = contact with prey or other predator and 1= maximum distance away. Analysis from all trials combibned, except for column marked * in which the effect is specific to Follow trials indicates significance in Follow trials only. There were no significant findings for Hunt trials alone.

| Direction to prey | Direction to other | Distance to prey | Direction to prey - Follow trials* |
| --- | --- | --- | --- |
| 15 ($t = -2.08, p = 0.048$)<br>- Frontopolar Area | 4 ($t = 2.43, p = 0.023$) -<br>Frontopolar area<br>22 ($t = -2.49, p = 0.020$) -<br>dLPFC | 18 ($t = -2.25, p = 0.035$) -<br>dLPFC | 3 ($t = -2.07, p = 0.049$) -<br>Frontopolar area |

### Dyadic Brain Synchrony

Before comparing INS during Hunt trials with Follow trials, we first examined INS in each trial type separately, comparing them to chance levels. Monte Carlo simulations were used to determine the alpha threshold of *p* < 0.180 in each dyad for significance, where we required that 4 dyads (k-threshold = 4) in each channel showed significance. In the high frequency range (0.1-0.2 Hz), channels 10, 13, and 15 of the bilateral frontopolar area exhibited significantly greater INS during Hunt trials compared to chance levels (Figure 5A) (*p* < .05). In contrast, during Follow trials, in the high frequency range (0.1-0.2 Hz), channels 7, 10, 13, and 15 of the bilateral frontopolar area showed significantly greater synchrony compared to chance levels (*p* < .05) (Figure 5B). In the middle (0.03-0.1 Hz) frequency range, channels 1, 7, 8, 9, 10, 13, 14, 15, 17, 18, 20, 22, and 23 of the bilateral dlPFC and frontopolar areas demonstrated significantly greater INS compared to chance levels (*p* < .05) (Figure 5B). In the low frequency range (0.02-0.03 Hz), channels 1, 8, and 20 of the left frontopolar area and right dlPFC demonstrated significantly greater INS compared to chance levels (*p* < .05) (Figure 5B). When we compared INS during Hunt trials with Follow trials, we found no significant differences in activity in any of the channels (*p* > .05).

**Figure 5.**
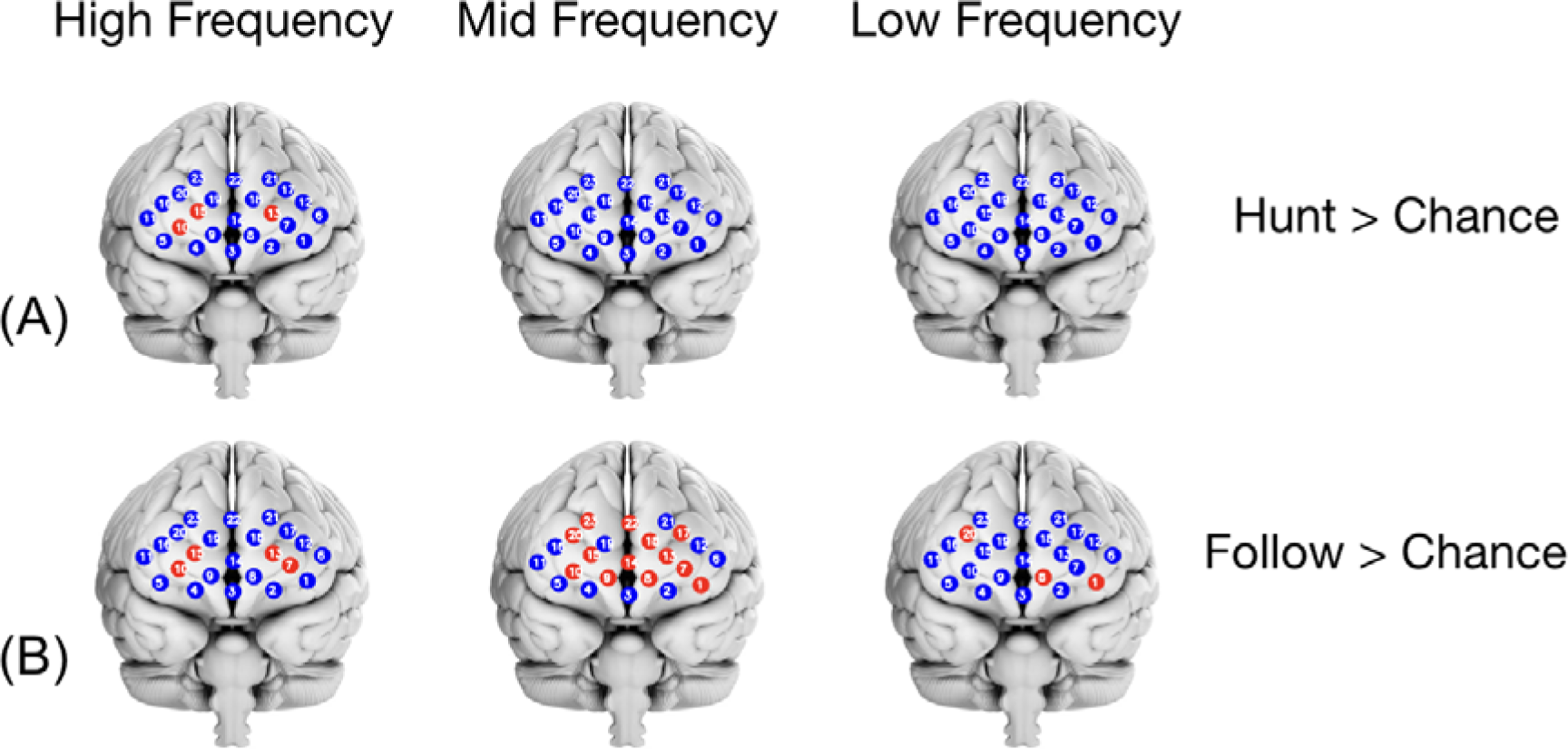
Interpersonal Neural Synchrony (INS) in Hunt > Chance and Follow > Chance contrasts. (A-B) Brain maps show anatomical locations of channels with significantly greater INS (red) during H unt and Follow trials vs Chance, across three frequency bands: low (0.02–0.03 Hz), middle (0.03–0.1 Hz), and high (0.1–0.2 Hz). Comparisons to chance reveal greater synchrony in both Hunt and Follow trials across the bilateral medial (frontopolar) and lateral PFC in all frequency bands. There were no significant findings for the Hunt > Follow or Follow > Hunt contrasts. Significant channels are highlighted in red.

We next considered an exploratory analysis to examine how behavioural variables might be related to synchrony. Trial averages of distance to prey, heading direction to prey, heading direction to other, and speed synchrony were correlated with hunting success, which was the time-window over which synchrony was calculated. This means any significant effects might be related to the capacity to detect synchrony rather than the synchrony itself, e.g. a correlation between synchrony and larger distances to the prey might occur because the epoch over which synchrony is captured has more fNIRS signal measured. However, we found the distance between predators during a hunt was not significantly correlated with the time of the trial. Thus, we explored whether any channels would show increased synchrony when hunters were closer or further away (average distance in a trial as the measure). We found that the average distance between predators in a trial was not significantly associated with neural synchrony in any frequency band (*p* > .05).

Overall, we found INS is greater in a range of channels across the lateral and medial (frontopolar) PFC during both Hunting and Following compared with chance levels, but no differences between the trial types and no association with average distance between predators in a trial block.

## Discussion

We combined a virtual minecraft world with fNIRS hyperscanning to investigate the neural and behavioural dynamics of cooperative hunting in humans. We found that distance to prey and speed synchrony between predators were significantly correlated with hunting success (time to catch prey). While there were some correlations with brain activity and distance and direction to prey in hunting, these were not significant when correcting for multiple comparisons. Using hyperscanning, we found evidence of synchronised brain activity across several prefrontal regions during hunting and following. No regions showed more synchrony for hunting than following or vice versa, and there were no overall differences between hunting and following when the brain activity was compared across individuals.

In many species, coordinated pursuit enhances hunting success by outmaneuvering prey (Bailey et al., 2012; Klein et al., 2021; Samuni et al., 2018; Watts & Mitani, 2002). From an analysis of the behaviour we found that successful hunting was higher when predators maintained a closer Euclidean distance to the prey and were more synchronised in their speed. It is unsurprising staying close to the prey is more successful. Greater proximity to a prey will increase the probability of a successful strike. By contrast, speed synchrony is associated with success is a more novel finding. It provides a potentially new insight into what makes a pair of human hunters more successful. Synchronizing speed may work favorably when predators keep pace with the prey together. If one predator far outstrips the other, it may diminish the coordination between predators. Consistent with this, wolf packs have been found to maintain a certain distance, and thus speed, to encircle or intercept prey (Muro et al., 2011).

It seems plausible that predators staying close together might have led to better prey capture, in that they could both strike to capture the prey and monitor each other’s movement more effectively. We found no evidence that proximity between predators was linked to success. In our experiment the prey was faster running in a straight-line than the predators, and thus if the predators just simply moved together as a duo it would result in poor success. Spreading out, to outflank the prey was required for success as is the case with wolf packs and other species that hunt as groups (Muro et al., 2011). Working with larger distances between predators could in theory lead to success, and this was not the case, rather success appears to rely on variation in distance between predators and the prey in an optimal way.

Analysis of group hunting in a range of species, including killer whales, octopus-fish groups, schooling fish, and wild baboons, indicates that successful hunting can involve one individual taking the lead and others adapting their paths in response to that lead predators movement (Couzin, 2009; Couzin et al., 2005, 2011; Pitman & Durban, 2011; Sampaio et al., 2024; Strandburg-Peshkin et al., 2015). Such a strategy makes sense for humans when it is not possible to use language to communicate (Bailey et al., 2012). To explore this we used the methods developed by Wu et al (2025) which tests for the presence of ‘pull’ events, where one predator’s movement (the leader) draws the other to adapt (the following predator). We found no evidence for such dynamics in our data. Visual inspection of the paths of predators fits this pattern, with predators appearing to be somewhat independent of each other. It seems possible such leader-follower dynamics might evolve as longer time was provided for the participants to engage in hunting as a dyad, as this would be the case with the species examined before. It is also possible that with three or more predators a hierarchy would emerge, with a more experienced or successful leader taking control of the leader-follower dynamics.

Our finding of increased intersubject neural synchrony between PFC regions during hunting is, to our knowledge, the first evidence that brain regions show increased synchrony between co-hunters during group hunting. Such synchronised brain activity is consistent with other tasks in which dyads must cooperate to solve a joint problem (De Felice et al., 2024; 2025). We found that synchrony was significantly above the range predicted by our null models in frontopolar PFC channels and at higher frequencies examined with fNIRS (0.1-0.2Hz). Synchrony at higher frequencies is consistent with the rapid behavioural dynamics required with group hunting. Increased synchrony in frontopolar channels provides evidence to developing models of spatial behaviour that seek to explain the neural systems involved in navigation and hunting behaviour in cluttered terrain (Evdardrsen et al., 2019; Goodroe and Spiers, 2022).

To explore the specificity of our findings we included a control task: following another person through the environment. Watching participants perform hunting and following, they appear very similar, both involving moving a person over space as a dyad. There were no differences in the variation in the distance and direction to the person-followed/prey between hunting and following trials. Notably, we found no differences in intersubject neural synchrony between hunting and following and frontopolar channels were also significantly more synchronous compared to a null model at high frequencies during the follow task. Thus, while there may be different cognitive demands in hunting and following, such differences do not manifest in our neural synchrony measures. The synchrony we observed in frontopolar channels in the hunting task and follow task could be considered a general demand to monitor a moving person in an environment than the specific demands of coordinating as a team for successful hunting, such as coordinating behavior to outflank the prey.

In the follow task we also found increased synchrony across a wide range of channels at lower frequencies. It is currently unclear what underlies these increases. The follow task had participants follow as exactly as possible the path of the predator, and this necessitated more aligned paths in the environment than hunting where out-flanking was evident. Whether this synchrony in movements is what is related to the broader range of channels significantly more synchronised than a null model in follow trials requires further investigation. Past research with fMRI has shown that when participants re-trace the paths of visually watch agents in a virtual world there is increased functional connectivity between the left lateral PFC and the medial temporal lobe (Wagner et al., 2023). Intracranial EEG recordings show theta oscillations linked to tracking the movements of another person in a real-world environment (Stangl et al., 2020). Replicating our task in fMRI and intracranial EEG would help test whether such patterns occur also in our follow and hunting tasks.

Past research has found brain activity correlated with the distance and/or direction to fixed goals during memory-guided navigation (Spiers and Maguire, 2007a; Viard et al., 2011; Sherill et al., 2013; Howard et al., 2014; Chadwick et al., 2015; Spiers et al., 2018; Shine and Wolbers, 2019; Patai et al., 2019; Basu et al., 2022). It has been speculated that these correlates relate to computation of the distance and direction to the goal during navigation (Spiers and Gilbert, 2015; Spiers and Barry, 2015; Patai and Spiers, 2021; Nyberg et al. 2022) and that such computations would extend to hunting prey where the distance and direction to the moving goal is coded (Goodroe and Spiers, 2022). While we observed activity in a left frontal channel was correlated with distance to the goal during hunting, this was not significant when we corrected for multiple comparisons. It may also be the case that the coding of moving goals operates through different neural mechanisms to fixed goals. Alternatively, we may have lacked sufficient measurements to detect the correlations as fNIRS can be worn comfortably for less time than fMRI recordings can be made and the depth of fNIRS recordings may have missed the key brain regions involved in such goal/prey tracking. Past research has suggested medial temporal lobe and deeper PFC regions are involved in coding the position of other mobile agents (Rodent, bat recordings, Yoo et al., Fine et al., Wagner et al, 2023; Stagl et al., 2020; Chericoni et al., 2026) and thus our channels may not have been able to detect activity in regions correlated with distance to moving prey.

### Limitations

While our study provides novel insights into the behaviour and possible neurobiological mechanisms involved in multi-person tracking of a moving human target, several limitations should be considered. As we noted previously, fNIRS was chosen as a method to explore interindividual neural synchrony, but it has limited depth for brain measurement, this prevented us from examining deeper brain regions, such as the dorsal anterior cingulate cortex and the hippocampus, both of which are critical in goal-tracking in humans and animals (Yoo et al., 2021; Stangl et al., 2020; Epstein et al., 2017; Patai and Spiers, 2021; Chericoni et al. 2026). Due to the time constraints on how long fNIRS can be worn comfortably we were limited in the conditions we could include and our power to detect correlations with distance and direction to the prey. Including solo hunting would have been useful to assess the specifics of group-hunting, which could be included in future fMRI research. While tasks in VR environments can show correlations with real-world behaviour (Bonavita et al., 2021; Coutrot et al., 2019; Goodroe et al., 2025), it is important to acknowledge that different behaviour may occur in real-world environments, in particular due to wider peripheral vision in the real-world and physical fatigue from running after prey. Methods to compare real-world to VR and immersive VR will be important to explore this (Zisch et al., 2024; Vigliocco et al., 2024).

### Future directions

This study is the first to explore dynamic goal tracking in a multi-agent setting using fNIRS hyperscanning, and it leverages Minecraft as an immersive platform to investigate neural dynamics in humans during goal-directed behaviour. This approach allows for the study of brain activity in a rich, interactive environment, which simulates complex, real-world behaviours, making it an ideal model for understanding collective goal tracking. Future studies could benefit from larger sample sizes to explore behaviour. Past research has shown the value of massive online experiments to explore world-wide demographics on behaviour (Allen et al. 2024; Coutrot et al., 2018, 2022a; 2025; Spiers Coutrot & Hornberger, 2023). The paradigm created here could be adapted into such a large-scale online test, allowing for systematic examination of group hunting behaviour in relation to the environmental features (Yesitepe et al., 2023), participants background growing up (Coutrot et al., 2022b; Yavuz & Spiers, 2026) and group sizes (Sridhar et al., 2023). It could also systematically help examine the impact of video games experience on hunting behaviour, which has been shown for virtual navigation (Connors et al., 2014; McLaren-Gradinaru et al., 2023; Yavuz et al., 2024; Yavuz & Spiers, 2026). Using a similar paradigm in fMRI and EEG would allow for more insights into the brain dynamics involved in group hunting. Simulating the predators and prey with reinforcement learning methods may be a particularly fruitful direction, where a more mechanistic interpretation of the hunting process could be described and understood and has been applied to navigation (de Cothi et al., 2022; Russek et al., 2017; Simon and Daw 2011; Zalabak et al., 2025; Zhang et al., 2026). Such models will benefit from considerations in how detours and the shape of terrain can systematically distort cognitive estimates of the distance, time, or layout of a space (Brunec et al., 2017; Zisch et al., 2024; Bellmund et al., 2020; Duncan et al., 2025). Finally, the study of neuropsychological patients would be useful to understand the brain regions contributing to hunting behaviour. Recent research has shown remarkably preserved navigation performance of hippocampal amnesics (Kolarik et al., 2016; Psiadian et al., 2024). It would be useful to determine if such patients would succeed in group hunting where simulating future trajectories seems important and the hippocampus is implicated (Hassabis et al., 2007; Bendor and Spiers, 2016; Olafsdottir et al., 2015). Patients with ventromedial PFC damage may also struggle to keep track of the prey, given problems with wayfinding linked with damage to this region (Ciaramelli, 2008; Spiers, 2008).

## Conclusion

This study highlights the role of the prefrontal cortex (PFC) in dynamic, group-based goal tracking during hunting behaviour. In a virtual group hunting task, we found greater intersubject neural synchrony during Hunt trials compared with chance levels across the lateral and medial (frontpolar) PFC, demonstrating the first evidence for neural synchrony during human group hunting. These results extend the animal literature, which has focused primarily on the behavioural aspects of hunting, to humans, by revealing both neural and behavioural dynamics associated with group hunting.

## Acknowledgements

We would like to thank all the participants who volunteered for this research, as well as the staff at the Institute of Sports, Exercise, and Health Research Centre at UCL for providing the facilities needed to conduct this study, including access to fNIRS technology and testing rooms. We would also like to thank Ada Nielsen for her valuable assistance with the Minecraft environment, as well as the Shockbyte and Minecraft support teams for their help with technical difficulties encountered with the Minecraft server and game. We thank Zita Patai for advice on analysis advice. This research was generously funded by the Leverhulme Trust.

## Data and code availability

The data and code used in this analysis is accessible at: https://osf.io/6wk34/

## Appendix

**Table A1.** Design matrices indicating the inclusion of regressors in each analysis model, categorized by task-based, parametric, and non-parametric variables. This table outlines which regressors are included in each of the design matrices (M1-M6). Each column corresponds to a different design matrix, and the presence or absence of each regressor is indicated across the rows. P = parametric regressor, T = task-based regressor and NP = non-parametric regressor. All behavioural metric regressors in all design matrices were convolved with the hemodynamic response function (HRF) to capture the temporal dynamics of brain activation, allowing us to examine their relationship with brain activity during the hunting task. In contrast, the physiological metrics (heart rate and breathing rate) were included in all design matrices in their raw, unconvolved form.

| Variable | M1 | M2 | M3 | M4 | M5 | M6 |
| --- | --- | --- | --- | --- | --- | --- |
| Hunt: Heading direction from the self to the prey<br>Hunt trials (NP) | * |  | * |  |  |  |
| Follow: Heading direction from the self to the prey<br>Hunt trials (NP) | * |  | * |  |  |  |
| Hunt: Euclidean distance to the prey (NP) | * |  |  | * |  |  |
| Follow: Euclidean distance to the prey (NP) | * |  |  | * |  |  |
| Hunt: Heading direction from the self to<br>prey*Euclidean distance to the prey (NP) |  |  |  |  | * |  |
| Follow: Heading direction from the self to<br>prey*Euclidean distance to the prey (NP) |  |  |  |  | * |  |
| Hunt trials (I) |  | * |  |  |  |  |
| Follow trials (I) |  | * |  |  |  |  |
| Hunt: Speed synchrony between the self and other<br>predator (T) |  |  |  |  |  | * |
| Follow: Speed synchrony between the self and other<br>predator (T) |  |  |  |  |  | * |
| Heart rate (NP) | * | * | * | * | * | * |
| Breathing rate (NP) | * | * | * | * | * | * |
| Nuisance regressor (in-between trials) (I) | * | * | * | * | * | * |
| Time (NP) | * | * | * | * | * | * |

**Figure S1.**
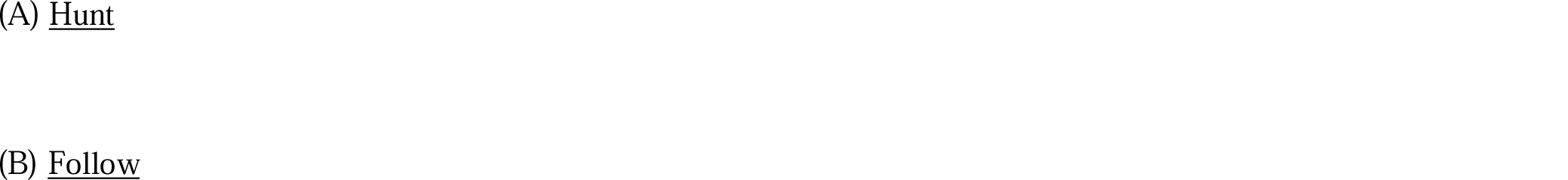
Example trajectories of the predators and prey during Hunt and Follow trials. **(A)** Example video showing a top-down view of the trajectories of the predators and prey as they unfold in real-time during a Hunt trial. **(B).** Example video showing a top-down view of the trajectories of the predators and prey as they unfold in real-time during a Follow trial.

